# Preparation of next-generation DNA sequencing libraries from ultra-low amounts of input DNA: Application to single-molecule, real-time (SMRT) sequencing on the Pacific Biosciences RS II

**DOI:** 10.1101/003566

**Authors:** Castle Raley, David Munroe, Kristie Jones, Yu-Chih Tsai, Yan Guo, Bao Tran, Sujatha Gowda, Jennifer L. Troyer, Daniel R Soppet, Claudia Stewart, Robert Stephens, Jack Chen, TF Skelly, Cheryl Heiner, Jonas Korlach, Dwight Nissley

## Abstract

We have developed and validated an amplification-free method for generating DNA sequencing libraries from very low amounts of input DNA (500 picograms – 20 nanograms) for singlemolecule sequencing on the Pacific Biosciences (PacBio) RS II sequencer. The common challenge of high input requirements for single-molecule sequencing is overcome by using a carrier DNA in conjunction with optimized sequencing preparation conditions and re-use of the MagBead-bound complex. Here we describe how this method can be used to produce sequencing yields comparable to those generated from standard input amounts, but by using 1000-fold less starting material.

## Introduction

In just the last few years, the development of second-generation sequencing (SGS) and third-generation sequencing (TGS) platforms, and the applications they enable, has driven the development of genomics, fundamentally altered our approach to life and medical sciences, and made possible the promise of personalized healthcare. ^(1, 3, 4)^ Library preparation for SGS and TGS can be labor-intensive and often requires starting material in the microgram range. ^(5, 6)^ These protocols require such large amounts of starting material due to the high rates of template loss during the wash steps that follow enzymatic reactions, with only a small fraction of the original starting material being represented in the final sequencer-ready product. This requirement for relatively large amounts of starting material can be a significant impediment to the sequencing of samples with limited amounts of DNA such as needle biopsy material, forensic or ChIP-seq samples, microorganisms refractory to growth in synthetic media, or when searching for rare sequence variants in unamplified nucleic acid samples. This is especially true when preparing unamplified libraries for single-molecule sequencing using the PacBio RS II sequencer. Unlike all SGS technologies, which rely on PCR and/or clonal amplification of DNA to generate thousands of copies of each template molecule for sequencing, PacBio library preparation does not require amplification of the DNA template during library preparation.^(7)^

A previously described method utilized the PacBio RS sequencer for direct sequencing from as little as one nanogram of input DNA.^(2)^ The method employs the use of random hexamer primers to anneal to the template DNA to provide the binding sites for the PacBio polymerase, thereby bypassing library preparation altogether. The method was applied to sequencing of ssDNA and dsDNA small genomes, with sequencing yields of mapped reads from less than a hundred to a few thousand per SMRT Cell. With the decrease in time and cost associated with library preparation, this method may be well-suited for rapid identification of infectious disease agents. However, because the sequencing yield produced is only a few thousand reads per SMRT Cell, application to larger or more complex genomes may be limited. Here we describe a simple, amplification-free method capable of producing standard sequencing yields by utilizing closed-circular plasmid DNA as a ‘carrier’ to minimize sample loss during library preparation. We hypothesized that closed-circular plasmid DNA will not receive the SMRTbell adaptor molecules that provide a priming site for the sequencing reaction, thereby mitigating loss of target DNA without contributing significantly to the sequencing output (Picture 1). The plasmid carriers are inexpensive and can be prepared in bulk. In addition to employing the use of a plasmid carrier during library construction, we have optimized the conditions of the final preparation steps of the libraries for sequencing, including sequencing primer annealing, polymerase binding, and MagBead binding. To maximize potential sequencing yield, we also reuse the MagBead-bound complex in subsequent sequencing runs. With the use of a circular plasmid carrier, optimized library preparation conditions, and re-use of the MagBead-bound complex, we demonstrate this method is capable of producing comparable, unbiased, per-SMRTcell sequencing yields from 1000-fold less starting material compared to the standard PacBio library preparation protocols.

**Picture 1.**
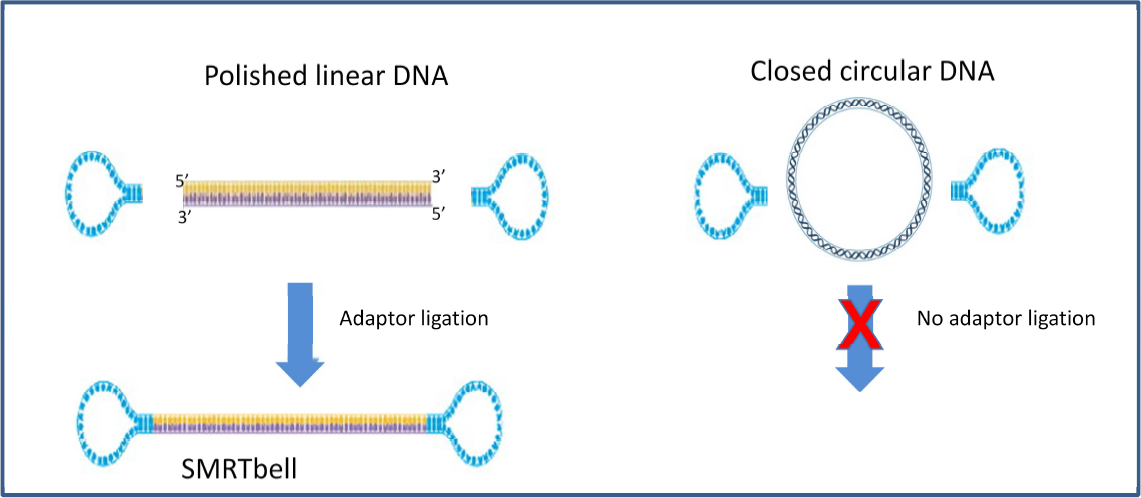
Principle of the low-input library preparation method. SMRTbell adaptors will ligate to linear DNA inserts of interest, but not to closed-circular plasmid DNA that is added as a carrier to the sample.

## Materials and Methods

The 2kb Low-Input Template Preparation and Sequencing protocol can be found on the Pacific Biosciences website in the Shared Protocols section of the SMRT Community Sample Network (SampleNet). λ genomic DNA (catalog no. 25250-010) was from Invitrogen™. HB101 *E. coli* genomic DNA was purified using the GenElute™ Bacterial Genomic DNA Kit (catalog no. NA2110) from Sigma-Alrich^®^. pUC18 plasmid (catalog no. 3218) was from Clontech. pBR322 plasmid (catalog no. N3033S) was from New England Biolabs®. Exonuclease III and Exonuclease VII kits (catalog no. EX4405K and EN510100, respectively) were from Epicentre®. All cleanup steps were performed using Agencourt AMPure XP beads (catalog no. A63881). A Covaris M220 Focused-ultrasonicator™ was used to shear DNA into 2kb fragments. Covaris g-Tubes™ were used with an Eppendorf® 5424 centrifuge to shear DNA into 20kb fragments. Pacific Biosciences’ DNA Template Prep Kits 2.0 (250bp - <3kb and 3kb – 10kb), (catalog nos. 001-540-726 and 001-540-835, respectively) were used to prepare the 2kb and 20kb fragment libraries, the DNA/Polymerase Binding Kit XL 1.0 and DNA/Polymerase Binding Kit P4 (catalog nos.100-150-800 and 100-236-500, respectively) were used during the annealing and binding reactions. A Blue Pippin™ from Sage Sciences was used to select large fragments in the 20kb fragment libraries. A GeneAmp® PCR System 9700 thermal cycler from Applied Biosystems was used for the annealing and binding reactions. Non-standard adjustments were made to the Annealing and Binding Calculator (versions 1.3.3, 2.0.1.0, and 2.0.1.2) provided by Pacific Biosciences to calculate the sequencing primer annealing, polymerase binding, and MagBead binding concentrations such that all library material available was loaded onto the sample plate for sequencing. The PacBio DNA Sequencing Kit 2.0 (catalog no. 001-554-002), SMRT Cell 8Pac v3 (catalog no. 100-171-800), MagBead Kit (catalog no.100-133-600), and MagBead Station were used for all sequencing. All 2kb sequencing was performed using C2 Chemistry and the XL polymerase on an RS I, with 2 x 55 minute movies, unless otherwise noted. Stage start was not enabled to maximize CCS yield. All 20kb sequencing was performed using the P4 polymerase on an RS II, with 1 x 120 minute movies. Stage start was selected to maximize insert read length. Sequence analysis was performed with SMRT portal, SMRT pipe, and SMRT View, versions 1.4, 2.0, and 2.1, all from Pacific Biosciences.

**See Supplemental Material for Bulk Plasmid Carrier Preparation, Low-Input Shearing, Library Preparation, Binding Calculator Adjustments and Sequencing Details**

## Results and discussion

Meeting PacBio sample input requirements can sometimes be a challenge and the sole limiting factor for several potential sequencing applications. If standard input amounts are used for library construction and a 20% recovery is assumed, the resulting final libraries contain tens of billions of SMRTbell molecules. This enables the possibility of sequencing hundreds of SMRT Cells when MagBead loading is employed, since only tens of millions of SMRTbell molecules are actually added to the SMRT Cell during sequencing. Considered in terms of mass, less than one nanogram is actually needed for sequencing one SMRT Cell (Supplemental Table 1). For a 2kb library, the number of SMRTbell molecules needed to produce standard sequencing yield are present in approximately 180 picograms. Theoretically, it should be possible to begin library construction with 5 nanograms and assume that 1 nanogram of final library resulting from a 20% recovery could still produce the standard expected sequencing yield. However, we have found that once the input amount used for library construction goes below 50 nanograms, the sequencing yield is significantly lower than the expected yield produced when using standard input amounts (Fig. 1). This may be due to decreased efficiencies in the enzymatic reactions throughout library construction as well as decreased AMPure bead binding efficiency at lower molar concentrations. The latter could also explain the lower recovery percentages seen when shearing small amounts of DNA (see Low Input Shearing section).

**Figure 1.**
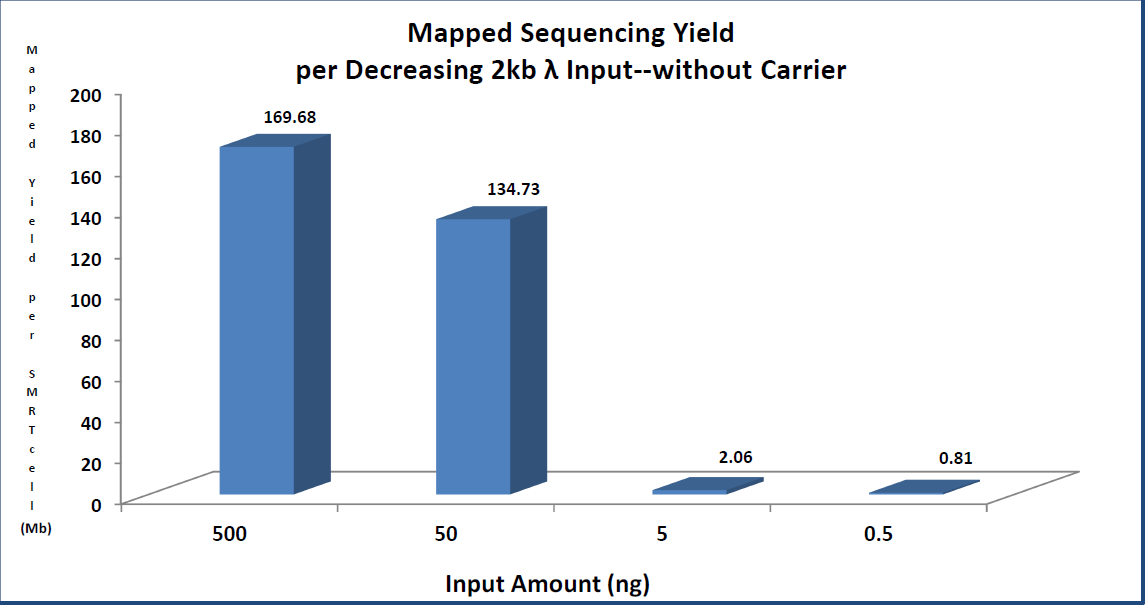
Mapped yield summary from decreasing amounts of 2kb λ DNA libraries without plasmid carrier.

Template loss during library preparation should be largely random and not sequence-specific. To determine if this is indeed the case, both 500bp and 2kb libraries were prepared by diluting decreasing amounts of sheared plasmid DNA (pBR322) into an amount of sheared lambda DNA that brought the total amounts up to 250 nanograms and 500 nanograms, respectively, which are the standard input requirements for 500bp and 2kb libraries. Libraries were prepared and subsequently sequenced. As expected, the number of reads sequenced from either the sheared plasmid or the sheared lambda was in direct proportion to that of the input amounts, by mass, during the dilution (Supplemental Fig. 1a and b). To further validate this finding, a serial dilution of one known 2kb amplicon into a second known 2kb amplicon was performed, beginning with a 10% spike-in by molarity and then serial diluting six times down to 0.15625%. Libraries were then prepared from each serial dilution using 500 nanograms as input for each and were subsequently sequenced. Again, a very strong correlation between expected and observed yield showed that there is no sequence-specific template loss during library preparation nor preferential sequencing bias (Supplemental Fig. 1c).

Because the end-repair and adaptor ligation reactions should be agnostic to carrier or target DNA, using linear DNA as a carrier provides a benefit of enabling quality control checks throughout library construction. However, sequencing yield of the low-input target will be limited by the molar ratio of target molecules to carrier molecules. As shown in Figure 1, a minimum of 50 nanograms is required to obtain standard expected sequencing yield, so for very low input target amounts (less than one nanogram), only approximately two percent of the sequencing yield could be expected to stem from the target compared to the carrier. For this reason, we decided to test the effectiveness of using a closed-circular carrier DNA molecule as a means to reduce the amount of carrier DNA sequenced, since the adaptors should not ligate compared to a linear carrier.

As described in the Library Preparation section, decreasing amounts of 2kb sheared lambda target DNA were used to prepare libraries, with 500 nanograms of the carrier pUC18 plasmid (2,686bp) spiked into each library after ligase inactivation. Input amounts for the target were chosen based on the expected recovery percentage as it relates to the resulting theoretical ratio of SMRTbell molecules to ZMWs (Supplemental Table 2). Sequencing was performed using C2 Chemistry and the XL polymerase on an RS I, with 2 x 55 minute movies. As shown in Figure 2, use of a circular carrier enables the sequencing of as little as 500 picograms of target DNA without an appreciable loss of mapped read throughput, generating as much as 160 Mb of mapped target DNA sequence in a single SMRT Cell run. Less than one percent of the overall mapped yield originated from the circular carrier, suggesting the exonuclease treatment is very efficient at digesting nicked plasmids to prevent undesired binding of the polymerase. As predicted in Supplemental Table 2, the mapped sequencing yields of the target DNA remained comparable to that of the control until the ratio of SMRTbells to ZMWs became limiting below 0.5 nanograms. As expected, there was very little coverage bias across the target genome (Supplemental Fig. 2), further supporting our assumption that template loss during library preparation and loading of the SMRTbells for sequencing is independent of sequence composition.

**Figure 2.**
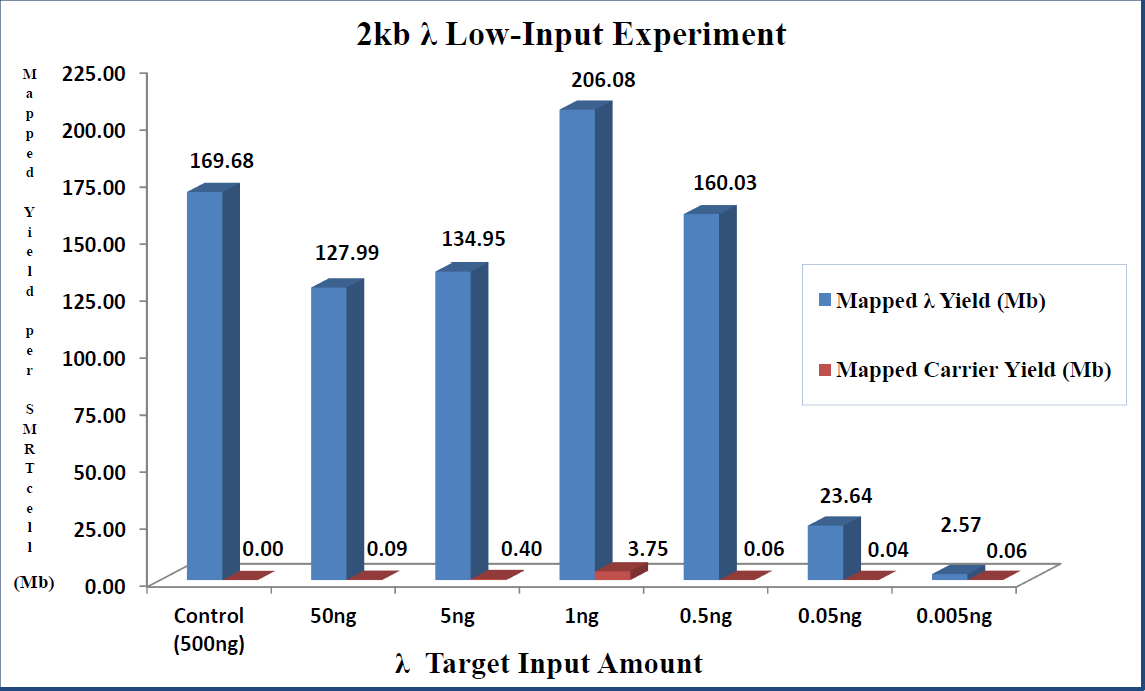
Mapped yield summary from decreasing amounts of 2kb λ DNA libraries in a plasmid carrier. 500ng of plasmid carrier (puc18) was used for each target input amount.

Much of the original PacBio MagBead-bound sample that is prepared for sequencing is left over in the bottom of the sample plate well following the initial run. In an attempt to use as much of the low-input target as possible for sequencing, PacBio Bead Binding Buffer was added in sufficient volume to accommodate the required dead volume that was lost from the first run, and samples were sequenced a second time.

While there was a noticeable decline in yield, the results were promising enough to warrant a third attempt at sequencing by again adding PacBio Bead Binding Buffer to the sample well and re-running (Fig. 3). Although the mapped yield from the two lowest target input amounts (5 and 50 picograms, respectively) was not comparable to those from the standard 500 nanogram input, re-sequencing the samples two more times produced approximately 5-50 million additional mapped bases, which could provide enough data depending on the goals of the sequencing study.

**Figure 3.**
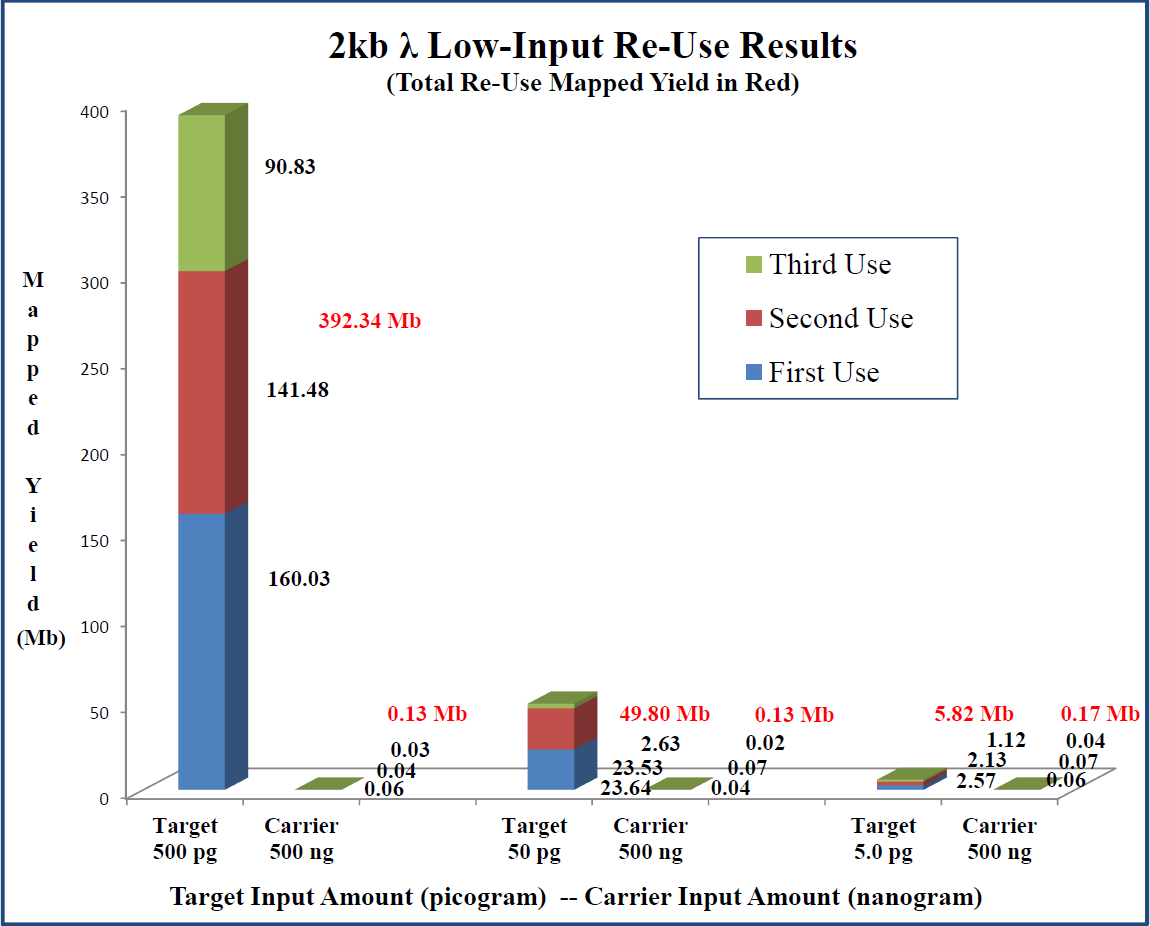
Mapped Yield summary from re-use of decreasing amounts of 2kb λ DNA libraries with a plasmid carrier.

Application of the 2kb low-input method to actual study samples produced sequencing yields comparable to those from the λ proof-of-principle experiments. Eight amplicons with very little starting material were sequenced using the 2kb low-input protocol (Fig. 4).

**Figure 4.**
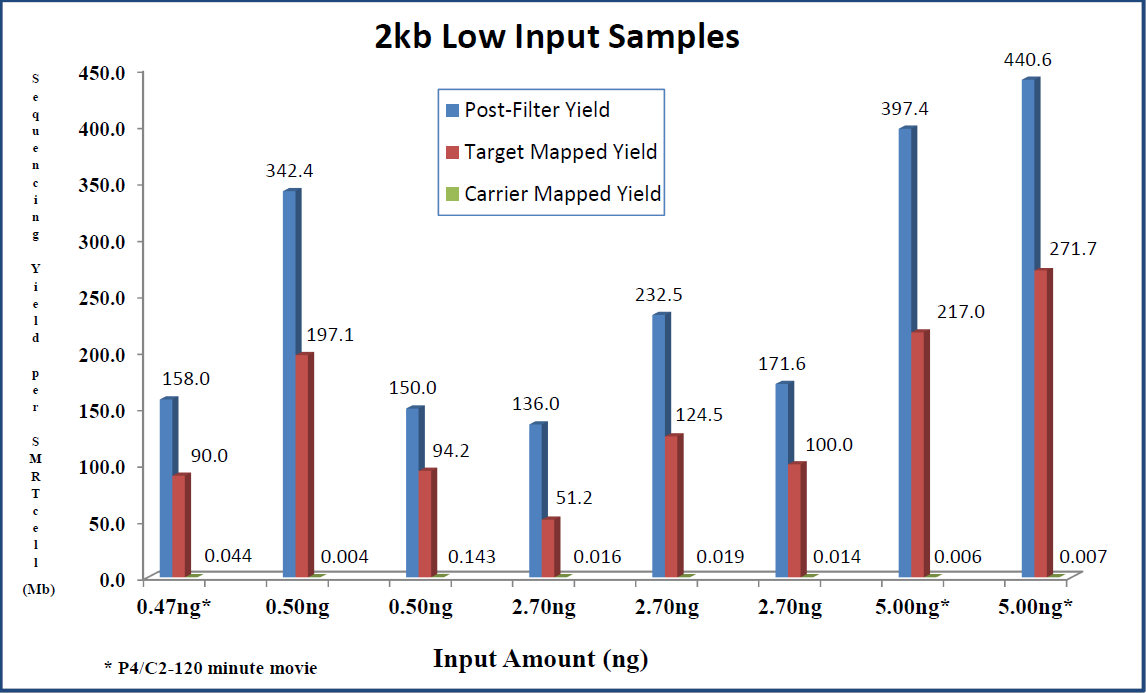
Sequencing yield summary from “real world” samples using the 2kb low-input protocol.

Input amounts varied from approximately 500 picograms to 5 nanograms. While mapped yields were lower for some samples compared to those from the λ experiments, the depth of coverage and number of circular consensus sequences produced for each sample was sufficient for determining haplotype phasing of highly homologous isoforms and detection of very rare quasispecies in a diversely mixed population of sequences (Supplemental Table 3).

To test the low-input protocol for long-insert libraries, we prepared 20kb libraries using *E.coli* as the target and pBR322 as the plasmid carrier as described in the Materials and Methods section and Supplemental Material. Samples were re-sequenced using the same re-use strategy employed in the 2kb low-input experiments. Figure 5 shows the post-filter and mapped yield for the *E. coli* target and plasmid carrier. The yields show a linear relationship with decreasing input amounts. Mapped yield from the plasmid carrier is much less compared to the 2kb experiments, presumably because most of the plasmid was removed by the Blue Pippin during size selection. Supplemental Table 4 summarizes the results from aggregating the data for each input amount and running PacBio’s Hierarchical Genome Assembly Process, version 2 (HGAP2). Coverage plots of the contigs from each input amount were fairly uniform and contained no drop-outs, with 100% of all bases being called (Supplemental Fig. 3). Consensus concordance was greater than 99.95 for all input amounts, demonstrating the utility of the low input protocol for small genome assembly even from 5 nanograms of starting material.

**Figure 5.**
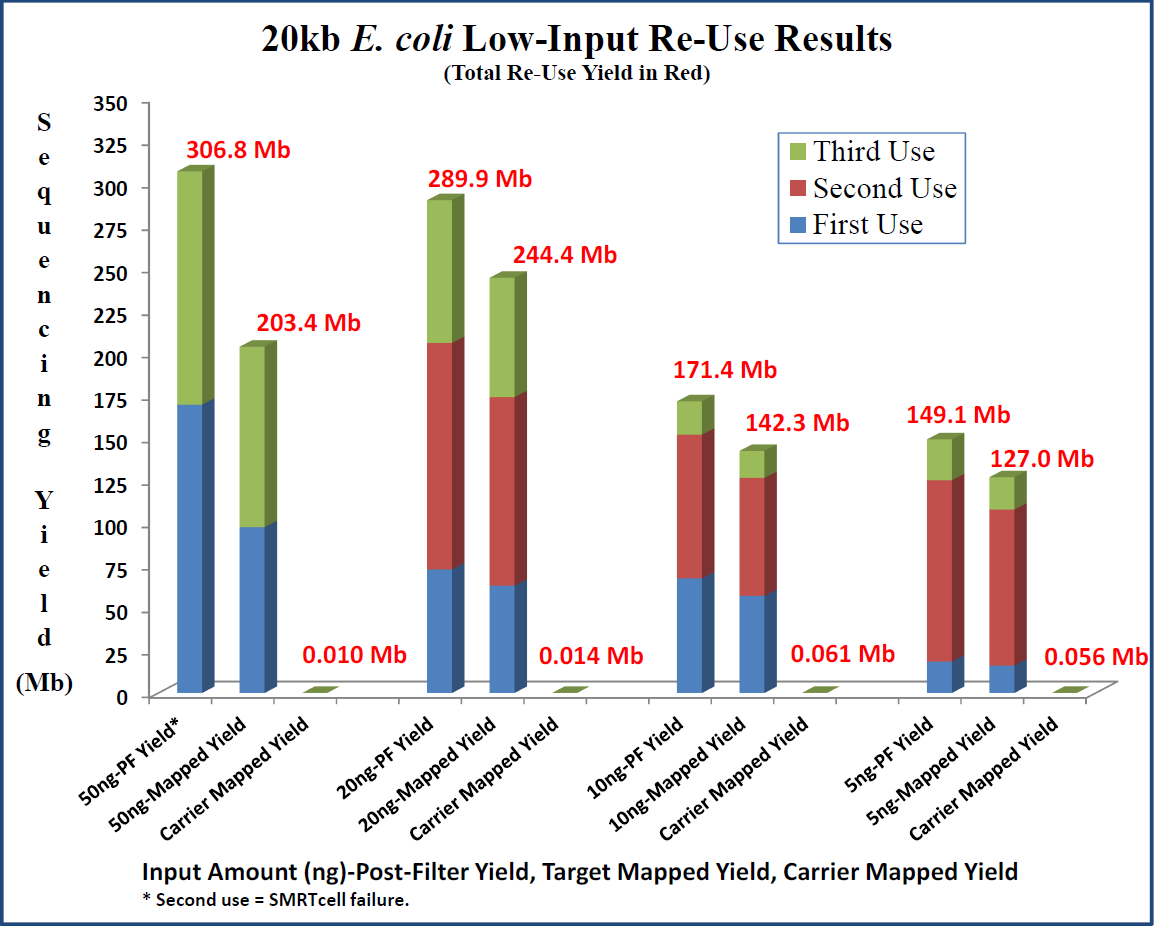
Sequencing yield summary from 20kb low-input experiments using decreasing amounts of *E.coli* target in plasmid carrier.

The methods described here demonstrate that it is possible to produce per-SMRT Cell sequencing yields comparable to both standard 2kb and 20kb libraries, using 1000-fold less starting material. The added cost and time from preparing the circular plasmid carrier is only a small fraction of the overall cost of library preparation, and can be performed in bulk to save time. While shearing and cleaning small amounts of starting material presently remains a challenge, there is still much room for optimization of current processes and development of new technologies to increase recovery rates. Projects for which the starting material is pure and high quality, but the amount available is the limiting factor, may now be enabled by the use of these methods.

## Competing Interests Statement

Leidos Biomedical Research, Inc. employees declare no competing interests.

PacBio Authors are full-time employees at Pacific Biosciences, a company commercializing single molecule, real-time sequencing technologies.

## Acknowledgments

We would like to thank PacBio for their input and guidance during optimization of the sequencing conditions.

